# Dynamic profile of malondialdehyde in renal and hepatic ischemia reperfusion injury: an explorative study of internal historical samples

**DOI:** 10.64898/2026.04.14.718142

**Authors:** Lene Devos, Tom Vanden Berghe, Diethard Monbaliu, Ina Jochmans

## Abstract

**Background:** Ferroptosis has emerged as a promising therapeutic target in IRI. However, it remains largely unclear how and when this iron-dependent regulated cell death manifests during IRI. Therefore, we explored malondialdehyde (MDA), a byproduct of lipid peroxidation, and glutathione peroxidase 4 (GPX4), as a marker of redox capacity, in multiple IRI models. With this explorative study, we aimed to uncover MDA dynamics in renal and hepatic IRI, which could provide valuable insights for future internal studies.

**Methods:** Historical plasma and tissue samples from rat and porcine models of renal and hepatic IRI were selected based on varying conditions of ischemic injury, reperfusion and perfusion. MDA was measured using a colorimetric assay with N-methyl-2-phenylindole, methanol, acetonitrile and hydrochloric acid and quantified at 595 nm. GPX4 protein concentrations were investigated using standard western blotting.

**Results:** In rat clamping models, plasma MDA concentrations revealed no difference between control and IRI settings. However, an increasing trend could be observed in tissue samples after IRI. Similarly, a decrease in tissue GPX4 concentrations was observed after IRI. In porcine studies, MDA concentrations were increased during reperfusion of kidneys exposed to prolonged warm ischemia and livers exposed to short periods of cold ischemia. Dynamic preservation could attenuate MDA concentrations.

**Conclusion:** We found that MDA and GPX4 are affected within the first hours after reperfusion, stressing the need for early sampling in studies focusing on characterizing ferroptosis. Moreover, MDA dynamics during organ perfusion revealed an increased vulnerability of ischemic organs to lipid peroxidation and a potential protective effect of dynamic preservation. These preliminary results should be confirmed in studies focusing on ferroptosis characterization, as notable observations regarding sample age and storage conditions and experimental design limit the validity of this study.

## 1. Introduction

Organ transplantation has become the most effective treatment for acute organ failure and end-stage organ disease thanks to major advancements in the field.(1) However, several limitations remain. Yearly, thousands of patients are added to the waiting list, where the limiting factor for treatment is the supply of good quality donor organs.(2) Organ quality is generally dictated by donor type, ischemic time, age, and presence of pre-existing disease.(3) The use of “extended criteria” donor organs narrows the gap between demand and supply, but not without risk. These grafts are more susceptible to ischemia reperfusion injury (IRI), which increases the occurrence of primary non-function and delayed graft function after transplantation. IRI, a detrimental type of tissue injury that occurs when blood supply returns to hypoxic tissue, is therefore the leading cause of high patient morbidity and mortality after transplantation.(4)

Ferroptosis, an iron-dependent form of regulated cell death, is a dominant driver of IRI and seems to be an effective therapeutic target in this field.(5-7) Ferroptosis is characterized by an accumulation of intracellular redox-active iron, which disturbs lipid metabolism through induction of lipid peroxidation driven by oxidative stress and enzymatic reactions. This lipid peroxidation causes excessive build-up of toxic lipid hydroperoxides in cellular membranes, which can be quantified by the lipid peroxidation byproduct, malondialdehyde (MDA). Insufficient control of these lipid hydroperoxides leads to membrane disruption and cell death.(8)

While the role of ferroptosis has been widely established in IRI, it is still unclear when and how ferroptosis manifests during transplantation. Recent evidence has confirmed the presence of ferroptosis during static cold preservation and normothermic kidney preservation, suggesting that ferroptosis emerges as early as organ preservation.(9, 10)

In this explorative study of historical samples, we aimed to uncover the dynamic profile of MDA in different models of renal and hepatic IRI present in our lab. Various conditions of ischemic injury, reperfusion and perfusion were investigated. Additionally, a cellular repair mechanism of lipid peroxidation, represented by glutathione peroxidase 4 (GPX4), was used in support of MDA to provide insights on the redox capacity during IRI. Since these preliminary results are derived from historical experiments that primarily focused on clinical endpoints, we would like to disclose that the conclusions from this report are limited and should be confirmed in further research using appropriate an experimental design (timepoints, sample sizes, control groups and markers).

## 2. Materials and methods

### 2.1. Animals

Sprague Dawley rats (250g-300g; 8-10 weeks; male and female; Janvier Labs, France) were housed at the KU Leuven Animal facility in individually ventilated cages with 14h/10h light-dark cycle and food (ssniff® R/M-H, Soest, Germany) and water *ad libitum*.

TOPIG_TN70 pigs (30-50 kg; prepubescent; Tojapigs Escharm, Nijmegen, The Netherlands). The pigs were housed at the KU Leuven Animal facility in a conventional in a closed housing system, in contact with other animals. They received a maintenance diet (MPig-H, ssniff, Soest, Germany) and were fasted 12h fasting for 12h prior to the experiments. The pigs had *ad libitum* access to water at all times.

The experimental and animal care protocols were approved by the KU Leuven Animal Care Committee (P014/2011; P209/2017; P054/2020; DMIT160/2021; DMIT061/2022) and conducted in accordance with European guidelines.

### 2.2. Models of IRI

#### 2.2.1. Rat models

##### 2.2.1.1. Rat model of bilateral renal clamping

Before surgery, analgesia was provided by subcutaneous injections of ropivacaine (5 mg/kg, Naropin®, Aspen, Ireland) at the site of incision and buprenorphine (0.01 mg/kg, Vetergesic®, CEVA, Libourne, France) in the neck. Anesthesia was induced with 5% isoflurane (IsoVet®, Piramal Critical Care BV, The Netherlands) and maintained with 1.5-3%. Animals were placed on heating pads during the entire procedure. After incision, the abdominal cavity was exposed with retractors and the intestines were moved aside with a damp gauze to facilitate renal pedicle dissection. After dissection, warm ischemia (WI) was induced with microaneurysm clamps with minimal time difference between kidneys. After 60 minutes of WI, the clamps were removed to allow reperfusion. Animals were sacrificed at 6 h,24 h or 48 h (intraperitoneal injection of 60 mg/kg Dolethal® (Vetoquinol NV, Belgium)) post-reperfusion. Sham animals went through the same operative procedure, without clamping, and were sacrificed immediately at the end of the procedure. Blood was collected and spun down (10’, 4°C, 2880 g) for plasma. Kidneys were collected at each timepoint. Samples were snap frozen and stored at −80°C for further analysis.

Note that samples from the first historical study were collected from experiments that used the same methodology as described above, with varying WI times (0, 45 or 60 minutes). Only anesthesia protocols were different, (Ketamine/Xylazine mixture) as described elsewhere.(11) Some samples were also stored at −20°C instead of −80°C (**Figure S1a**). Moreover, at the time of analysis, the storage time was approximately ten years for the first study, whereas samples from the new study were analyzed within weeks after the experiment.

##### 2.2.1.2. Rat model of partial liver clamping

Rats were anesthetized using 5% isoflurane (IsoVet®, Piramal Critical Care BV, The Netherlands) for induction and 1.5-3% for maintenance. Animals were placed on heating pads during the entire procedure. Before the start of surgery, analgesia was provided by subcutaneous injections of ropivacaine (5 mg/kg, Naropin®, Aspen, Ireland) at the site of incision and buprenorphine (0.01 mg/kg, Vetergesic®, CEVA, Libourne, France) in the neck. After incision, the abdominal cavity was exposed by retractors and forceps were used to lift the sternum upwards. Using damp cotton tips, the portal triad was exposed and the left and median lobe (±70% of the liver) were clamped. After 60 minutes of WI, the atraumatic clamp was removed to allow reperfusion of the ischemic lobes, inducing severe, reversible injury. Sham animals went through the same operative procedure, without clamping. Longitudinal blood samples were taken at 6h (short isoflurane induction; tail vein; 250-500 µL) and 24h (sacrifice; intraperitoneal injection of 60 mg/kg Dolethal® (Vetoquinol NV, Belgium); abdominal bifurcation, 6-9 mL) of reperfusion in each rat. Samples were spun down (10’, 4°C, 2880 g) for plasma and were snap frozen and stored at −80°C until use. Tissue samples of all lobes were collected after 24h of reperfusion, snap frozen, and stored at −80°C until use. MDA analysis was performed within weeks after the experiment.

#### 2.2.2. Porcine models

##### 2.2.2.1. Porcine model of *ex situ* kidney preservation and transplantation

The primary results of this study have been published elsewhere, along with detailed information about the surgical procedure.(12, 13) After nephrectomy, the renal arteries were cannulated, and the kidneys were flushed with 250 mL of cold preservation solution (IGL-1 at 4 °C, Institut Georges Lopez, Lissieu, France). 500 mL of autologous blood was collected for normothermic whole blood reperfusion. Before mounting on the perfusion circuit, kidneys were rinsed with 200 mL of cold Ringer’s solution (4 °C). Three experimental groups were investigated: (no WI/CI (cold ischemia)) control kidneys that were mounted on the perfusion circuit without any ischemia time; (22h CI) cold ischemic kidneys (CI) were stored on ice in IGL-1 for 22 h; (60’WI) warm ischemic kidneys underwent 60 min of *in situ* WI through ligation of renal artery and vein before nephrectomy. Each group contained 3 kidneys. To mimic transplantation, kidneys underwent normothermic whole blood *ex situ* reperfusion for 4 h, during which perfusate samples were collected, centrifuged at 2880*g* for 10 min at 4°C, aliquoted, snap frozen, and stored at −80°C for further analysis. MDA analysis was performed one to three years after the experiment.

For the hypothermic oxygenated machine perfusion study, kidneys were placed on a perfusion machine after nephrectomy and perfused with cold University of Wisconsin Machine Perfusion Solution for 20 h, as previously described.(14) Prior to cold perfusion, kidneys were exposed to either 60 minutes of WI (60’WI) or no ischemia (no WI/CI). Afterwards, they underwent 4 h of normothermic whole blood *ex situ* reperfusion. Samples were collected at 15 minutes, and 1, 3, 6, 9, 18, 19 and 20 h of cold perfusion and processed as described above. The same samples were collected during normothermic whole blood *ex situ* reperfusion. MDA analysis was performed a few months after the experiment.

##### 2.2.2.2. Porcine model of *ex situ* liver preservation and transplantation

The primary results of these studies have been published elsewhere, along with detailed information about the surgical procedure.(15, 16) For the prolonged normothermic preservation model (15), the liver was prepared for hepatectomy through *in situ* cold flush with IGL-1 preservation solution (Institute George Lopez, Lissieu, France) and dissection. After hepatectomy, the liver was cold stored for approximately 2 h and prepared for perfusion. After vessel cannulation, the livers underwent 24 h of normothermic reperfusion with a red blood cell based perfusate. Perfusate samples were collected at 15 and 30 minutes, and 1, 2, 3, 6, 12, and 24 h, centrifuged at 2880g for 10 minutes at 4 °C, snap frozen, and stored at –80 °C until further analysis. MDA analysis was performed one to three years after the experiment.

For the normothermic machine perfusion and whole blood ex situ reperfusion model(16), livers were exposed to either 60 minutes of WI or not. After hepatectomy and 2 h of cold storage, livers were perfused with a red blood cell based perfusate for 6 h at normothermic conditions, followed by 12 h of normothermic whole blood *ex situ* reperfusion, mimicking the first hours after transplantation. Perfusate samples were collected at 15 and 30 minutes, and 1, 2, 3, 6 and 12 (when applicable), centrifuged at 2880g for 10 minutes at 4 °C, snap frozen, and stored at –80 °C until further analysis. MDA analysis was performed one to three years after the experiment.

### 2.3. Ferroptosis markers

#### 2.3.1. Colorimetric lipid peroxidation assay

MDA, a byproduct of lipid peroxidation, was quantified as previously described.(17) Briefly, 100 µL of tissue homogenate, plasma or perfusate was mixed with 325 µL reagent mixture containing 1-Methyl-2-phenylindole (3558-24-5, Merck, Germany), acetonitrile (75-05-08, Merck, Germany) and methanol (67-56-1, Merck, Germany). At 70°C, in the presence of 37% hydrochloric acid (75 µL; 258148, Sigma-Aldrich, USA), MDA reacts with 1-Methyl-2-phenylindole, producing a purple-blue colored adduct. Following centrifugation (10 minutes, 4°C, 15,000g), the absorbance of the supernatant was measured at 595 nm using an iMark™ microplate absorbance reader (Bio-Rad, USA). MDA concentration was calculated using a standard curve generated from 1,1,3,3-tetramethoxypropane (102-52-3, Acros Organics, Belgium) as the source of MDA. Tissue MDA measurements were corrected based on protein concentration (B6916, Bradford reagent, Merck, Germany) and expressed as µmol/g.

#### 2.3.2. Glutathione peroxidase 4

GPX4 concentration was determined by western blotting as previously described.(11) Briefly, 50 µg of tissue homogenates were prepared in Laemmli Sample buffer (2X (161-0737); 4X (161-0747), Bio-Rad, USA) containing β-mercaptoethanol (3 minutes at 95°C) and loaded on a precast gell (#4569036, Mini-PROTEAN TGX gells, Bio-Rad, USA). SDS-PAGE was performed to separate proteins based on kD (150 V). Next, proteins were blotted on a membrane (#1704156, Transfer Turbo Transfer Pack, Bio-Rad, USA), that ultimately was blocked (room temperature, 1h, 5% milk powder), incubated with primary antibody (4°C, overnight), washed (PBS-T) and incubated with secondary horseradish peroxidase coupled antibody (room temperature, 45 minutes). After washing (PBS-T), immunoreactive bands were visualized through chemiluminescence. Band intensity was quantified using Chemidoc MP techniology and associated Imagelab software (Bio-Rad, USA).

### 2.4. Statistical analysis

Detailed statistical analysis and data representations can be found in each figure caption. A p-value<0.05 was considered statistical significant. All analyses were perfomed in GraphPad Prism 10.6.1.

## 3. Results

### 3.1. *In vivo* investigation of malondialdehyde and glutathione peroxidase 4 during reperfusion

#### 3.1.1. Bilateral renal clamping experiments in rats

In the first study, MDA was investigated in plasma and tissue samples at different timepoints (0h, 1h, 3h, 6h, 24h and 48h of reperfusion) from historical female controls that underwent sham operations, 45 minutes of WI or 60 minutes of WI. Noticeable differences were observed between plasma samples stored at −20°C and −80°C, suggesting that storage conditions have a tremendous effect on MDA concentrations and that lipid peroxidation continues after sampling (**Figure S1a**). Based on these observations and the extended age of these samples (∼10 years), MDA concentrations in these samples are likely not reliable and are, therefore, not reported. Nevertheless, an overview of plasma and tissue concentrations can be found in **Figure S1b-d**.

In a second part, samples from a new experiment, including sham controls and animals exposed to 60 minutes of WI (60’WI), were investigated for both MDA concentrations. Both plasma and tissue MDA concentration in male and female animals showed no difference over time compared to sham animals (p=0.192, p=0.409, respectively)(**Figure 1a**). A slight increasing trend in tissue could be observed in male animals after 60 minutes of warm ischemia, whereas MDA concentrations female animals seem to recover towards 48h of reperfusion. This trend is similar to the creatinine profile in this model (**Figure S2a**,**b**). In support to MDA, GPX4 was investigated as an indication of the kidney redox capacity. In both male and female animals, a significant change could be observed over time in both male and female animals (p=0.043). A slight reduction in GPX4 concentrations could be observed after 6 h of reperfusion in animals exposed to 60 minutes of WI compared to sham controls (**Figure 1c, Figure S1b**), corresponding to the peak of injury in these animals (**Figure S1d**,**e**). However, GPX4 expression recovers during reperfusion, exceeding sham concentrations after 48h of reperfusion in some cases.

**Figure 1.**
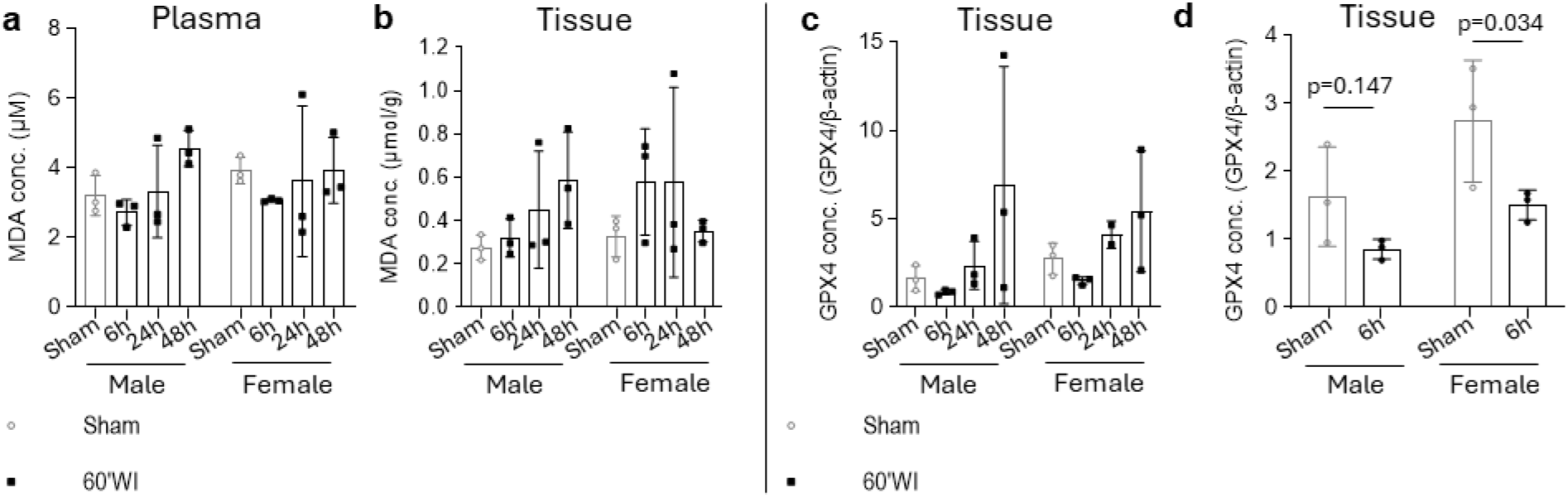
MDA and GPX4 concentrations over time in bilateral renal clamping experiments in rats. (**a**) Plasma MDA concentrations in rats exposed to 60 minutes of warm ischemia or sham operations at different times of reperfusion. (**b**) Kidney tissue MDA concentration in rats exposed to 60 minutes of warm ischemia or sham operations at different times of reperfusion. (**c**) Kidney tissue GPX4 concentration in rats exposed to 60 minutes of warm ischemia or sham operations at different times of reperfusion. (**d**) Bar/dotplots representing GPX4 concentrations in sham animals and animals exposed to 60’WI and 6h of reperfusion. (**a-d**) Groups consisted of both male (n=3/group) and female (n=3/group) rats. Each dot represents an individual rat. Statistical analysis and data representation: (**a-b**) Two-way ANOVA with Dunnet’s multiple comparison (mean + SD) (**c**) Two-way ANOVA with Tukey’s multiple comparison (mean + SD); (**d**) Two-way ANOVA with Fisher’s LSD multiple comparison (mean + SD); Non-significant p-values were not included in the graphs.

#### 3.1.2. Partial liver clamping experiments in rats

After 24h of reperfusion higher MDA concentrations could be observed in females exposed to 60 minutes of WI (60’WI) in both left (p<0.001) and median (p=0.038) lobe compared to males. In the left lobe, this was also significantly higher compared to sham female animals (p=0.009) (**Figure 2a**,**b**). In male animals, no significant change could be observed compared to shams. Also, plasma MDA concentrations were similar between the groups (**Figure 2c**). At 24h post-reperfusion, GPX4 concentrations were significantly lower in female median lobes compared to shams (p=0.029), whereas no difference could be observed in male animals. In the left lobe, no differences were observed in male and female groups. **Figure S1f**,**g** displays the corresponding injury levels in this model, which peaks at 6h of reperfusion and significantly decreases at 24h post-reperfusion, suggesting that 24h post-reperfusion might be too late to observe meaningful differences between groups.

**Figure 2.**
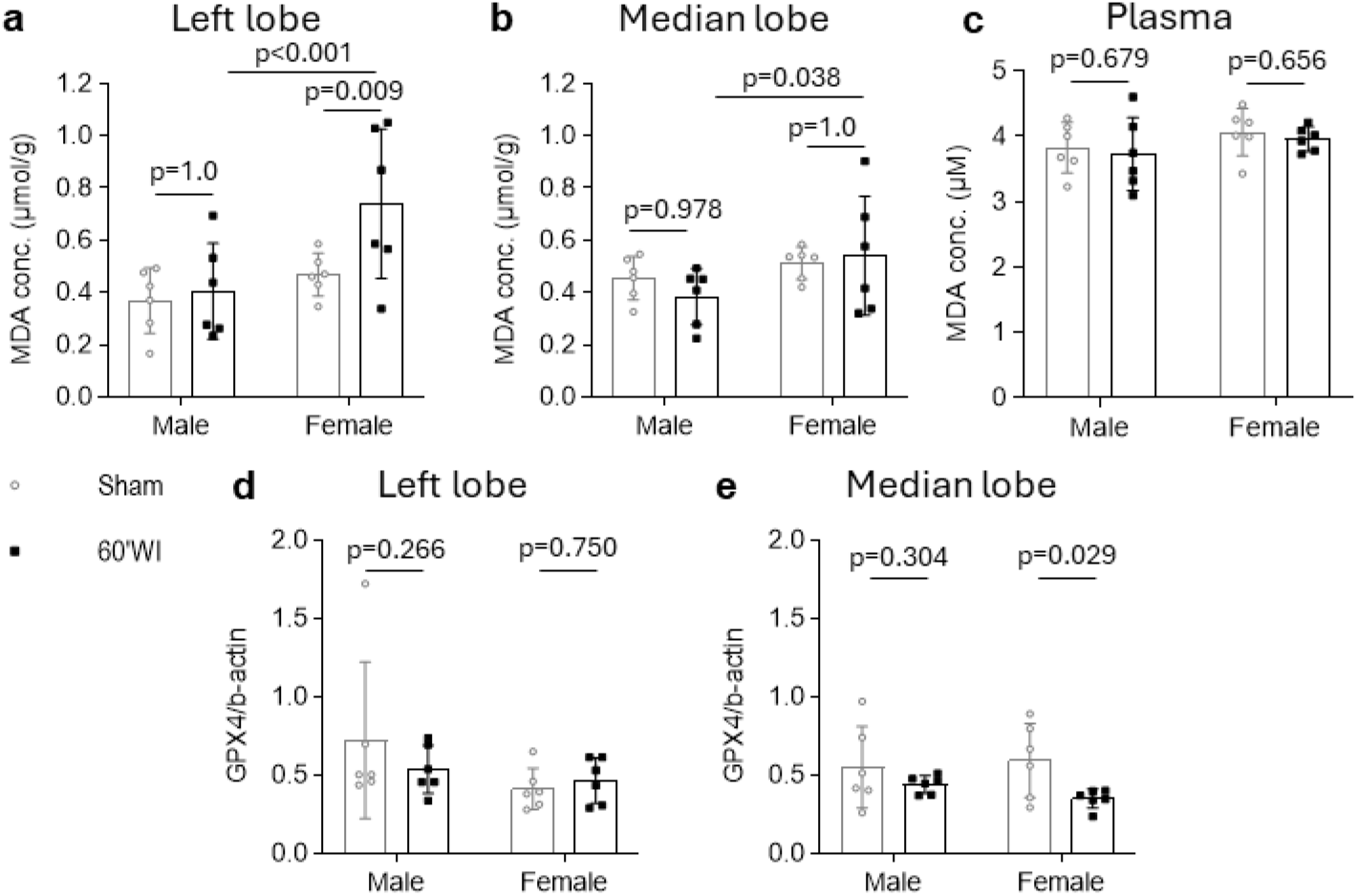
MDA and GPX4 concentrations in partial liver clamping experiment in rats. (**a**) Liver left lobe MDA concentrations after 60 minutes of warm ischemia and 24h of reperfusion compared to sham operated animals. (**b**) Liver median lobe MDA concentrations after 60 minutes of warm ischemia and 24h of reperfusion compared to sham operated animals. (**c**) Plasma MDA concentrations in rats exposed to 60 minutes of warm ischemia and 24h of reperfusion compared to sham operated animals. (**d**) Liver left lobe GPX4 concentration after 60 minutes of warm ischemia and 24h of reperfusion compared to sham operated animals. (**e**) Liver median lobe GPX4 concentration after 60 minutes of warm ischemia and 24h of reperfusion compared to sham operated animals. Both male and female animals were included. Each dot represents an individual rat (n=7/group). Statistical analysis and data representation: (**a-c**) Two-way ANOVA with Tukey’s multiple comparison (mean + SD); (**d-e**) Two-way ANOVA with Fisher’s LSD multiple comparison (mean + SD)

### 3.2. *Ex situ* dynamics of malondialdehyde during reperfusion in different donor types and preservation methods

#### 3.2.1. Porcine kidney preservation and reperfusion

In perfusate samples of pig kidneys exposed to different types of ischemia, a significant increase over time could be measured in the group exposed to 60 minutes of WI (60’WI, p=0.033) (**Figure 3a**). Interestingly, no significant change could be observed in perfusate from kidneys that underwent no ischemia (no WI/CI, p=0.634) and 22h of CI (22h CI, p=0.284) (**Figure 3a**), suggesting that kidneys exposed to prolonged WI are more prone to undergo lipid peroxidation during reperfusion. During hypothermic oxygenated perfusion, both uninjured and 60’WI kidneys showed a significant increase in perfusate MDA concentration (no WI/CI: p=0.016; 60’WI: p=0.036). However, no significant change could be observed during whole blood reperfusion (no WI/CI: p=0.453; 60’WI: p=0.439) (**Figure 3b,c**). Moreover, overall concentrations during reperfusion remained below 5 µM when exposed to hypothermic perfusion, whereas MDA concentrations without prior exposure to hypothermic perfusion rose up until 10 µM in kidneys exposed to 60 minutes of WI, proposing a protective effect of hypothermic oxygenated machine perfusion against lipid peroxidation during reperfusion.

**Figure 3.**
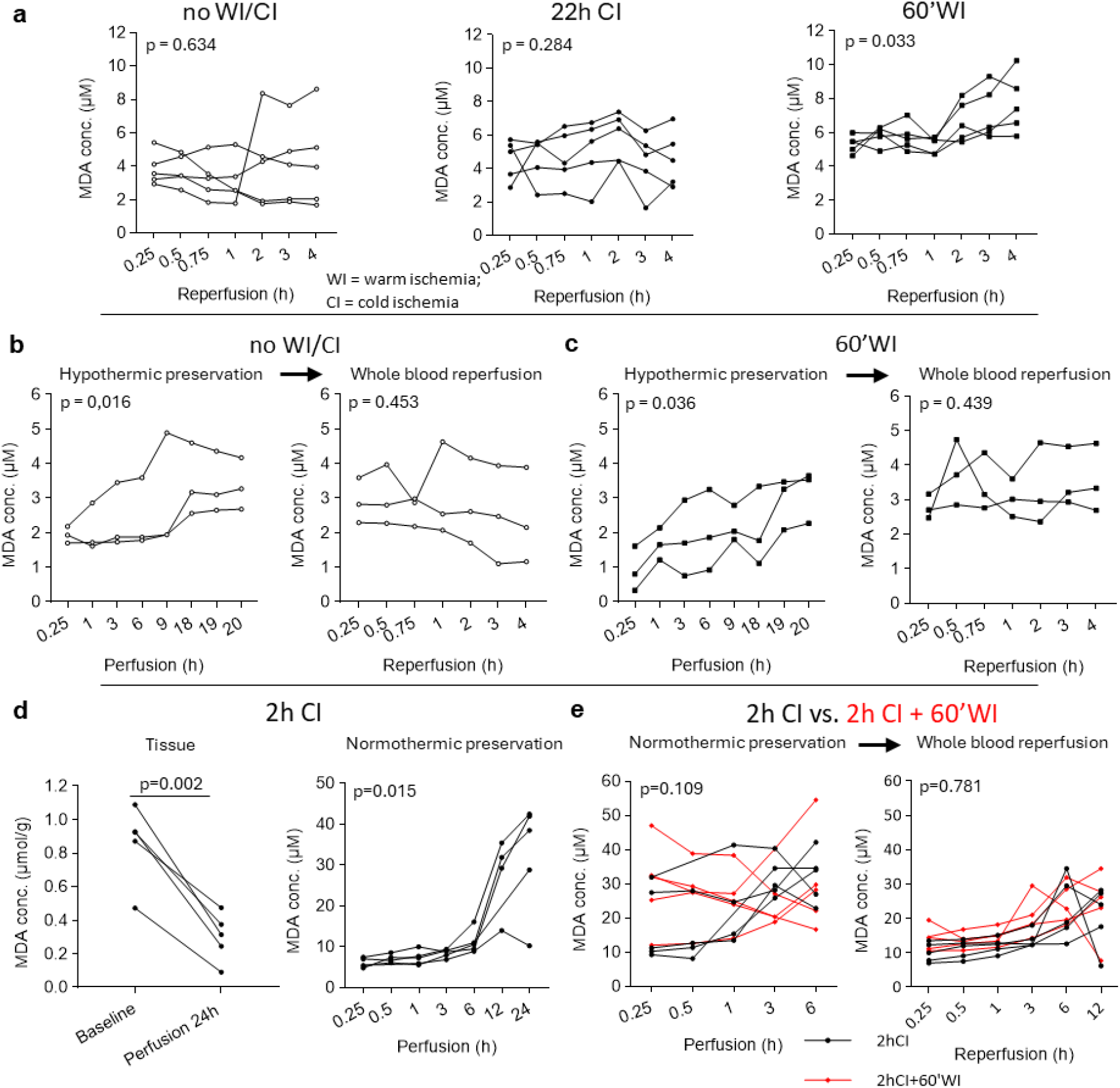
MDA concentrations during *ex situ* liver and kidney preservation and transplantation. (**a**) Perfusate MDA concentrations during normothermic whole blood reperfusion of kidneys exposed to different ischemia times (no WI/CI, 22h CI and 60’WI, n=3/group). (**b**) Perfusate MDA concentrations of kidneys exposed to no ischemia (no WI/CI, n=3), which underwent 20h of hypothermic oxygenated perfusion, followed by 4h of normothermic whole blood reperfusion. (**c**) Perfusate MDA concentrations of kidneys exposed to 60 minutes of warm ischemia (60’WI, n=3), which underwent 20h of hypothermic oxygenated perfusion, followed by 4h of normothermic whole blood reperfusion. (**d**) MDA concentration of baseline tissue samples and samples after 24h of normothermic perfusion of livers exposed to 2h CI (left) and perfusate MDA concentration of livers exposed to 2h CI and 24h of normothermic perfusion. (**e**) Perfusate MDA concentrations of livers exposed to 2h CI, which underwent 6h of normothermic perfusion, followed by 12h of normothermic whole blood reperfusion. Statistical analysis and data representation: (a) Repeated measures One-way ANOVA (all, mean + SD); (**b-c**) Friedman test (no WI/CI hypothermic preservation, median + 95% confidence interval), Repeated measures One-way ANOVA (others, mean + SD); (**d**) Respectively, paired t-test, mixed effects analysis (mean + SD); (**e**) Respectively, mixed effects analysis (mean + SD), two-way ANOVA (mean + SD)

#### 3.2.2. Porcine liver preservation and reperfusion

MDA concentrations in pig liver tissue seemed to decline over time during prolonged normothermic perfusion (p=0.002), whereas perfusate concentrations increased over time (p=0.015)(**Figure 3d**). Interestingly, a major burst of MDA could be observed after 6h of perfusion (**Figure 3d**), demonstrating that long-term perfusion is accompanied with increased lipid peroxidation. In a second experiment short-term normothermic preservation of livers showed high MDA concentrations over time in both livers exposed to 2h CI or 2h CI and 60 minutes of WI, but no significant increase over time could be measured, nor a significant difference between both groups (p=0.109) (**Figure 3e**). Also no difference between both groups was observed in the subsequent whole blood reperfusion phase (p=0.938) (**Figure 3e**). Despite increasing MDA concentrations during whole blood reperfusion, short-term normothermic preservation of the livers did seem to cause a milder increase during *ex situ* reperfusion (**Figure 3e**).

## 4. Discussion

This explorative study provides an overview of MDA dynamics, a surrogate marker of lipid peroxidation, in internal models of renal and hepatic IRI. Overall, the conclusions in this study are hampered by sample age, storage conditions, small sample sizes and the lack of adequate timepoints and control groups. A detailed discussion of these observations is provided below.

In the historical bilateral clamping experiments, we exposed the effect of different storage conditions on MDA, showing that MDA formation and, therefore, lipid peroxidation can proceed during storage, as demonstrated in previous studies. Addition of anti-oxidants, like butylated hydroxytoluene, and fast sample processing (snap freezing and storage at −80°C) can prevent this increase.(18-20) However, it is advised to analyze samples as soon as possible after experiments are conducted. Based on these observations, we decided not to discuss the results from overly aged samples, as their MDA concentrations are likely not reliable. We also advise to interpret the graphs in **Figure S1** with caution. In both *in vivo* settings of renal and hepatic IRI, plasma MDA concentrations did not reveal remarkable differences between sham and WI animals. This might be explained by the high reactivity of MDA. MDA possibly reacts with various plasma proteins, forming adducts that can obscure the measurement of this marker.(21) Indeed, the colorimetric method used in this study cannot detect these adducts.(17) This reactive prolife of MDA underline the need for additional markers.(22) Therefore, we included GPX4 as a secondary, which has been linked to ferroptosis in both kidney and liver.(23, 24)

In a more recent study of renal IRI, MDA and GPX4 concentrations in tissue seemed to align with the extend of renal degradation (creatinine and AST, respectively) in this model. A slight increase in MDA could be observed, which extended until 48h of reperfusion in males. GPX4 concentrations, on the other hand, were decreased after 6h of reperfusion but recovered in later timepoints, which could point towards a compensation mechanism. Unfortunately, the small sample size in this study (n=3/group) limits the validity of these results.

In the liver partial clamping model, we mainly observed changes in tissue MDA and GPX4 concentrations of female animals. However, due to the study design, tissue samples were only available at 24h of reperfusion. At this timepoint, liver injury markers (AST and ALT) are largely reduced compared to the peak concentrations at 6h of reperfusion. Therefore, we could not provide a complete overview of the dynamics of these markers in relation to the injury observed in this model, which limits the conclusion from the current observation. We suspect that the inclusion of earlier timepoints, could have provided valuable insights, as shown in renal IRI.

In porcine studies, we found that kidneys exposed to prolonged WI show increased vulnerability to lipid peroxidation during reperfusion, which could be attenuated with hypothermic oxygenated perfusion. These findings correlate with the improved clinical outcomes linked to this preservation methods in kidney transplantation.(25, 26) Moreover, kidneys exposed to no ischemia or prolonged CI did not show a significant increase of lipid peroxidation over time, underlining the detrimental impact of prolonged WI in kidneys.(27) In contrast to kidneys, MDA concentrations increased over time in livers exposed to 2h CI, which corresponds to the known differences in tolerance towards CI times in kidney and liver.(28, 29) Moreover, similar to recent evidence in kidneys, we showed that prolonged normothermic preservation triggers a burst of lipid peroxidation in livers, which might be correlated with hemolysis.(9) However, after short-term normothermic perfusion, cold stored livers showed a milder MDA increase over time, suggesting that short-term NMP can dampen lipid peroxidation during reperfusion, also corresponding to its clinical performance.(30) Moreover, no difference could be observed with livers exposed to prolonged WI. However, this condition was not tested without exposure to NMP, which could have attenuated MDA concentrations in this group. These observations, suggest a promising potential of MDA as a predictive biomarker in *ex situ* perfusion settings, where MDA builds up in the circuit without rapid clearance, in contrast to *in vivo* settings.

Lastly, while we largely focus on MDA and GPX4 as a supportive marker, conclusions regarding ferroptosis remain limited. Characterization of ferroptosis is not achieved by a universal marker. Instead, multiple pathways should be investigated to characterize a ferroptosis signature. Brent R. Stockwell, head of the founding lab of ferroptosis, stated that markers of lipid peroxidation should be supported by at least three other markers detecting changes in gene expression, mitochondrial changes and/or transferrin receptor mobilizations.(22) Therefore, further research should be done in powered studies, including multiple markers, appropriate timepoints and controlled sample processing.

In summary, this study confirmed the effect of storage time and conditions on MDA concentrations in plasma and tissue samples, stressing the need for appropriate sample processing. Moreover, *in vivo* studies of renal and hepatic IRI uncovered the volatile nature of MDA and the need to include early timepoints of reperfusion and multiple markers when studying ferroptosis induced-IRI. In *ex situ* porcine settings of organ preservation and reperfusion, MDA analysis suggests increased vulnerability of ischemic organs to lipid peroxidation and a potential protective effect of dynamic preservation.

## 5. Acknowledgements

Special thanks to Tine Wylin for her technical support in sample processing and lab analyses of MDA and GPX4. Ina Jochmans is a Senior Clinical Investigator of the Research Foundation Flanders (FWO; 1800126N). Funding was provided by FWO SBO (S001522N), without involvement in study design, data collection, data-analysis, manuscript preparation and publication decisions.

## Notes

### Competing Interest Statement

The authors have declared no competing interest.

