## Supplemental Figures S1 and S2 for "Dynamic profile of malondialdehyde in renal and hepatic ischemia reperfusion injury: an explorative study of internal historical samples"

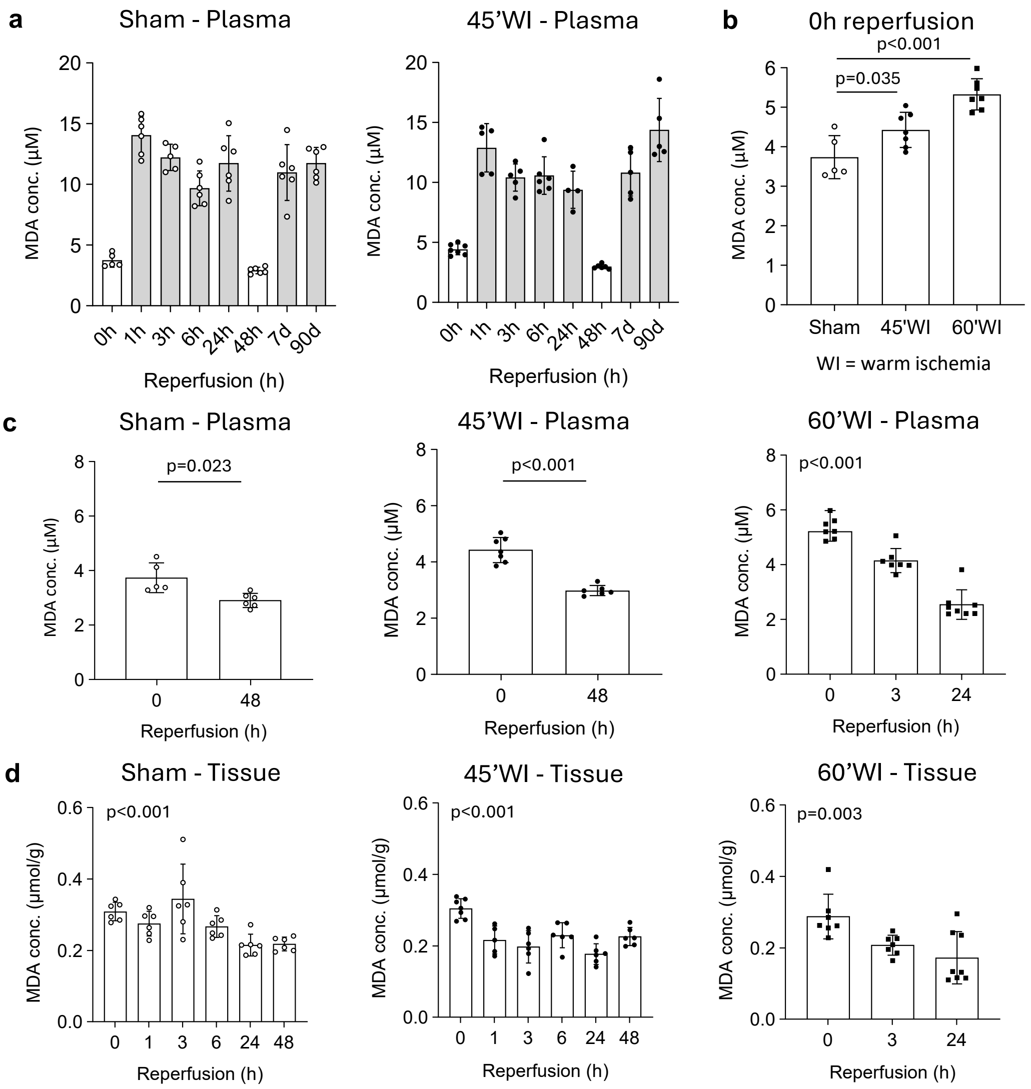


**Figure S1. MDA concentrations are affected by sample age and storage conditions.** (**a**) Bar/dotplots showing MDA concentrations over time in samples stored at different conditions (-20°C: gray bar; -80°C: white bar). (**b**) Bar/dotplots showing plasma MDA concentrations right after warm ischemia (when applicable) in sham, 45'WI and 60'WI groups. (**c**) Plasma MDA concentrations at different times of reperfusion in sham animals and animals exposed to 45 or 60 minutes of warm ischemia. (**d**) Kidney tissue MDA concentrations at different times of reperfusion in sham animals and animals exposed to 45 or 60 minutes of warm ischemia. (**a-d**) Groups included 5-8 female animals. Each dot represents an individual rat.

Statistical analysis and data representation: (**a**) No statistical analysis was performed (mean + SD); (**b**)One-way ANOVA (mean + SD); (**c**) Welch's t-test (sham+45'WI, mean + SD), Kruskal-Wallis test (60'WI, median + 95% confidence interval); (**d**) One-Way ANOVA (sham+45'WI, mean + SD), Kruskal-Wallis test (60'WI, median + 95% confidence interval);

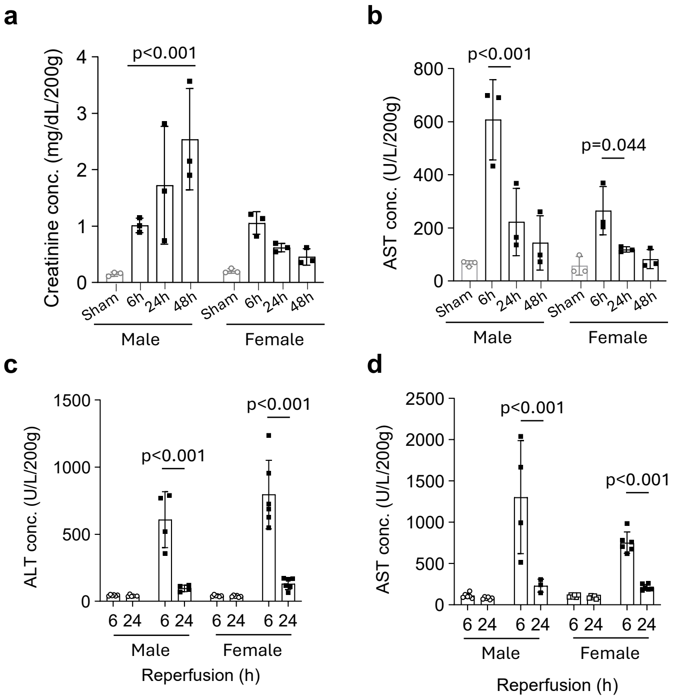


**Figure S2. Injury profile in renal and hepatic clamping models.** (**a**) Creatinine concentration in bilateral renal clamping model, as a marker of renal function. (**b**) Aspartate transaminase (AST) concentration in bilateral renal clamping model, as a marker of renal injury. (**c**) Alanine transaminase (ALT) concentration in partial liver clamping model, as a marker of liver injury. (**d**) AST concentration in partial liver clamping model, as a marker of liver injury. Statistical analysis and data representation: (**a-d**) Respectively; two-way ANOVA and mixed effects analysis with Tukey's multiple comparison (mean + SD). (**a,b**) Individual datapoint from separate animals. (**c,d**) These data represent repeated measures at 6h and 24h of reperfusion.
